# Multisensory expectations shape olfactory input to the brain

**DOI:** 10.1101/283242

**Authors:** Lindsey A. Czarnecki, Andrew H. Moberly, Cynthia D. Fast, Daniel J. Turkel, John P. McGann

## Abstract

The mammalian brain interprets sensory input based on prior multisensory knowledge of the external world, but it is unknown how this knowledge influences neural processing in individual sensory modalities. We found that GABAergic periglomerular interneuron populations in the olfactory bulb endogenously respond not only to odors but also to visual, auditory, and somatosensory stimuli in waking (but not anesthetized) mice. When these stimuli predict future odors, they evoke enhanced interneuron activity during the time odor normally occurs. When expectations are violated by omitting an expected “warning tone” before an odor, odor presentation evokes a burst of interneuron activity. The resulting GABA release presynaptically suppresses neurotransmitter release from the axon terminals of olfactory sensory neurons, the cells that transduce odor in the nasal epithelium and communicate this information to the brain. Expectations, even those evoked by cues in other sensory modalities, can thus affect the very first neurons in the olfactory system.

## Main Text

The brain learns statistical patterns in the stream of incoming sensory information, thus building a cognitive representation of the contingencies among stimuli that allows it to anticipate future events. Most real-world events engage multiple sensory modalities, producing learned stimulus “associations” in which stimuli in one modality can elicit expectations about likely impending occurrences in other modalities. Sensory stimuli that are congruent with these expectations are detected more readily^1^ and reacted to more quickly^2^, while *violating* expectations by changing or omitting an expected cue disrupts basic sensory processing, delays behavioral responses, and decreases task accuracy^2-8^. Physiological correlates of sensory expectations have principally been observed in the cerebral cortex^6^, but earlier sensory regions may also be influenced by expectations^9^, presumably through the numerous “top-down” projections from other brain regions. Here we use optical neurophysiology in a mouse model to demonstrate that in the mouse olfactory system, top-down cross-modal (non-olfactory) sensory information reaches all the way to the first neurons in the brain’s olfactory circuit and that establishing and violating expectations can modulate the synaptic terminals of primary sensory neurons at the input to the brain.

In the olfactory system, olfactory sensory neurons (OSNs) in the nasal epithelium transduce the odor into a neural signal, which passes down their axons in the olfactory nerve to glomeruli in the brain’s olfactory bulb. In these glomeruli the OSN axons drive projection neurons from the bulb to the olfactory cortex, but they also directly and indirectly drive a population of *Gad2*-expressing periglomerular (PG) interneurons^10^ (Fig. 1A). Note the distinctive exclusion of the Gad2-PG cell bodies from the glomeruli innervated by OSNs, but also that their dendritic and axonal processes distribute diffusely within and between glomeruli (Fig. 1B). These *Gad2*-PG cells release γ–aminobutyric acid (GABA) onto the OSN terminal both spontaneously and in response to odor stimulation. This combined tonic and feedback inhibition that can suppress OSN synaptic signaling to the brain via GABA_B_ receptors on their axon terminals^11-13^. *Gad2*-PG cells also receive extensive centrifugal input from other brain regions, including projections from cortical and neuromodulatory regions^14-23^. This convergence of peripheral and descending central inputs could uniquely position *Gad2*-PG cells to govern primary olfactory input to the brain on the basis of “top-down” information such as stimulus expectations.

**Fig. 1.**
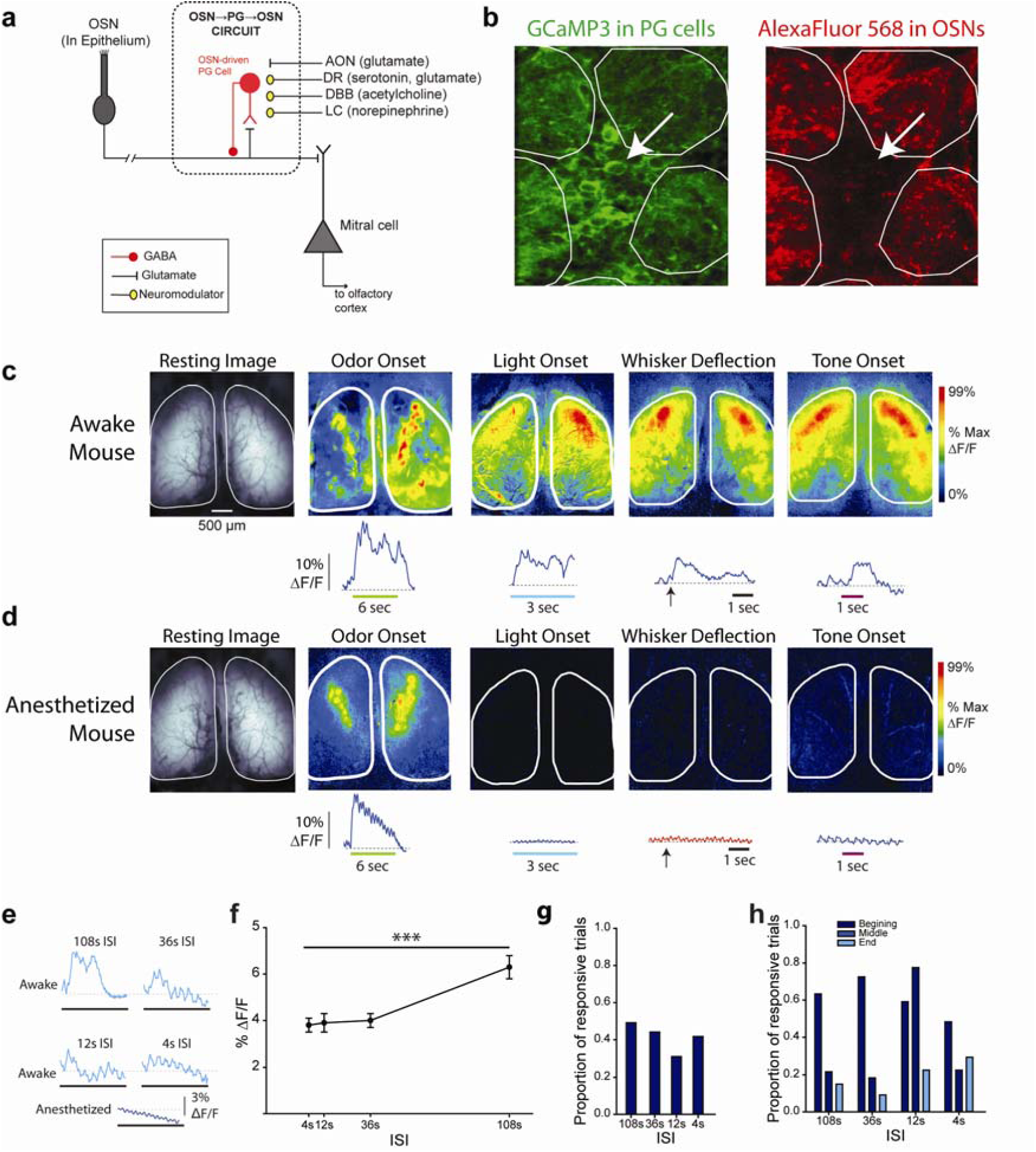
Olfactory bulb periglomerular interneurons populations receive multimodal sensory input. *A)* Schematic of OB circuitry showing convergence of olfactory sensory neuron (OSN) input with projections from higher brain areas in the OSN-periglomerular cell (PG) circuit. *B)* Fluorescence image of OSNs (red) anterogradely labeled from the nasal epithelium with surrounding Gad65-expressing PG interneurons (green). *C)* Representative pseudocolor maps and fluorescence traces showing stimulus-evoked calcium signals in GAD65-PG cells in awake mice. Colored bar or arrow notes stimulus presentation and duration. *D)* Only olfactory responses were observed while under anesthesia (same animals as in C). *E*) Single trial examples of light-evoked calcium signals in GAD65-PG cell populations averaged across the dorsal olfactory bulb at 108-, 36-, 12-, and 4-sec ISIs in an awake animal (light blue) or anesthetized animal (dark blue). *F*) Peak light-evoked response magnitude across ISIs (expressed mean ± SEM; 8 bulbs in 4 animals). *G*) Proportion of trials including light-evoked responses across ISIs. *H*) Distribution of light-evoked response peak latencies across the three one-second time bins composing each trial.

## Results

### Olfactory bulb interneurons respond to non-olfactory stimuli

Olfactory bulb glomeruli are the foundation of odor perception and coding, reflecting the identity and concentration of odors in the environment. As previously shown, in awake head fixed PG-GCaMP mice, which express the calcium indicator GCaMP^24^ in PG cells, odor presentations evoked strong calcium signals in odor-specific sets of glomeruli (Fig. 1C, Odor). Surprisingly, the population of PG cells sometimes also made stimulus-locked responses to the presentation of a visual stimulus (the bright 470 nm light emitted from the microscope during imaging; Fig. 1C, Light), a somatosensory stimulus (a brief deflection of the mouse’s whiskers; Fig. 1C, Whisker), and an auditory stimulus (a 1 kHz, 74 dB tone; Fig. 1C, Tone). Unlike odor-evoked activity, these cross-modal responses were not spatially localized to a specific subset of glomeruli and were spatially similar across stimuli. When mice were imaged both awake and under anesthesia, the calcium signals evoked by olfactory stimulation were qualitatively similar in both states. However, the responses to cross-modal stimulation (light, tone, and whisker brush) were completely eliminated by anesthesia (Fig. 1C vs. 1D).

In awake mice, repeated presentations of visual stimuli with a 108 sec interstimulus interval (ISI) evoked stimulus-locked responses on about half of the trials, while shorter ISIs evoked responses that were significantly smaller and less frequent (Fig. 1E-G). This contrasts with odor-evoked responses, which were very reliable (odors always evoked a response in experiments below) from trial to trial. In rodents, unexpected or novel stimuli typically evoke sudden inhalations, so we evaluated whether infrequent non-olfactory stimuli evoked exaggerated inhalations that might drive activity in the PG cell population in a subset of mice where we simultaneously monitored intranasal airflow with an implanted intranasal thermocouple during light presentation. We observed that in many cases light onset evoked both a change in sniffing and diffuse excitation of the PG cell population throughout the olfactory bulb (Fig. 2A_i_), but that there were also instances in which the PG response occurred without a corresponding change in sniffing (Fig. 2A_ii_) and instances in which neither clearly changed (Fig. 2A_iii_). To definitively rule out a causal role for sniffing, we thus repeated the experiment in two awake, tracheotomized mice and observed that while odor-evoked activity was completely eliminated in the absence of peripheral airflow (Fig. 2B, Odor), the response to cross-modal stimuli persisted (Fig. 2B, Light, Whisker, and Tone). These experiments indicate that the cross-modal signaling of PG cell populations throughout the olfactory bulb requires that the mouse be awake, but does not require peripheral input.

**Fig. 2.**
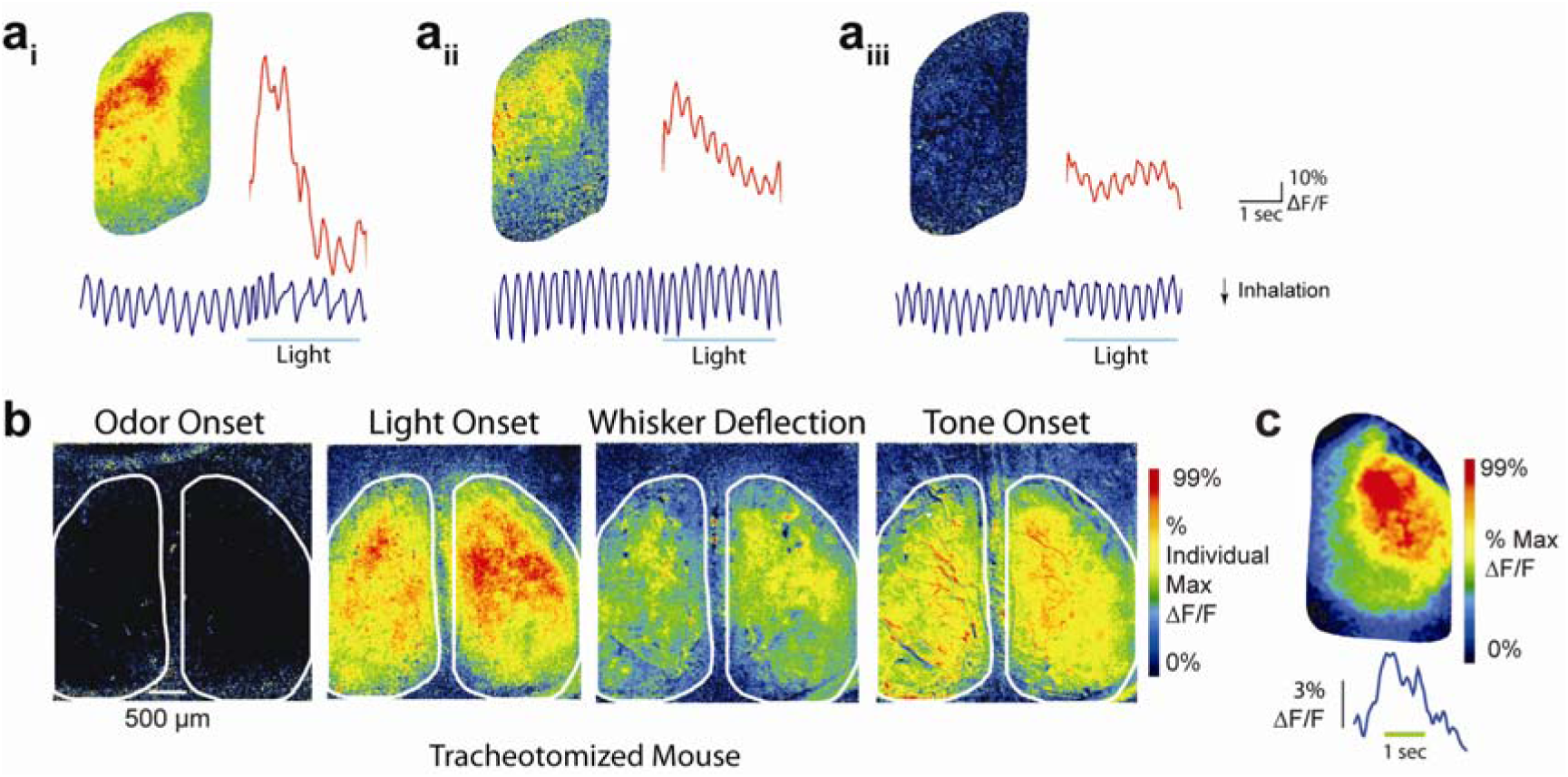
Cross-modal responses are not due to intranasal airflow. A) Representative pseudocolor maps with fluorescence traces (red) along with thermocouple respiration traces (blue) in response to light onset (examples from a PG-GCaMP3 reporter mouse). In some cases (Ai) respiration changes co-occur with changes in PG population activity. Other instances show no changes in respiration with clear increases in PG activity (Aii), or no changes in respiration with no obvious change in PG activity (Aiii). B) Representative pseudocolor maps across stimuli after tracheotomy scaled to individual maxima (example from a PG-GCaMP6f reporter mouse). C) Pseudocolored response map of light-evoked calcium activity in a mouse expressing GCaMP6f in PG cell populations via viral transfection into the glomerular layer.

Further control experiments confirmed that responses to cross-modal stimuli were observed in *Gad2-*GCaMP3 reporter mice (e.g. Fig. 1C-D; see Methods), *Gad2*-GCaMP6f reporter mice (e.g. Fig. 2B), and in mice expressing GCaMP6f in *Gad2-*PG cells through viral transfection (Fig. 2C). No such responses were observed in mice expressing GCaMP6f, synaptopHluorin, or Calcium Green in the OSN axon terminals interspersed among the *Gad2*-PG cells’ processes (data not shown; representative instance observable in Figure 4D_ii_ vs 4E_ii_). This rules out the possibility that these diffuse signals were optical artifacts of metabolic signaling or stimulus-evoked movement.

**Fig. 3.**
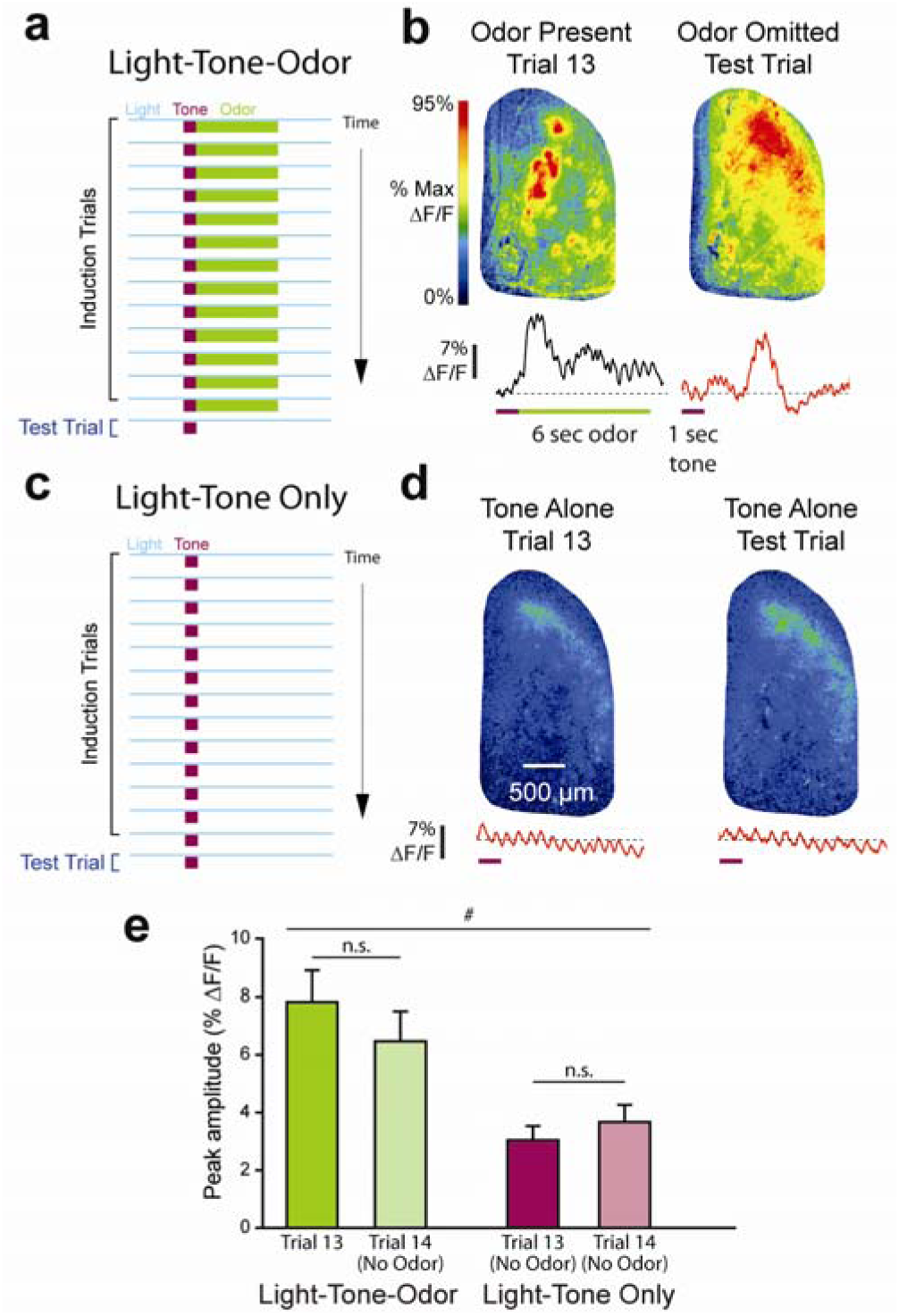
Expectation of odor presentation facilitates GABAergic signaling in the olfactory bulb. *A)* Paradigm schematic. *B)* Representative pseudocolor response maps of peak odor-evoked response on the final odor present trial, corresponding peak response on the following odor-omitted trial, and corresponding fluorescence traces. *C)* Control paradigm schematic. *D*) Pseudocolor response maps of peak response amplitude from the six seconds following tone presentation (corresponding to the odor presentation as in A) from the 13th and 14th trials (as in B) with corresponding fluorescence traces. Note that the response maps are scaled to the same absolute values as A to illustrate the difference in response amplitude. *E)* Light-tone-odor GCaMP3 peak signals were significantly larger than corresponding peak responses in the light-tone only experiment (mixed model ANOVA *F* = 9.3, *η*_*p*_^*2*^ = 0.48 from 12 mice, mean ± SE; # *p* < 0.02). Response amplitudes were not significantly different between the 13^th^ trial and 14^th^ trial in either paradigm.

**Fig. 4.**
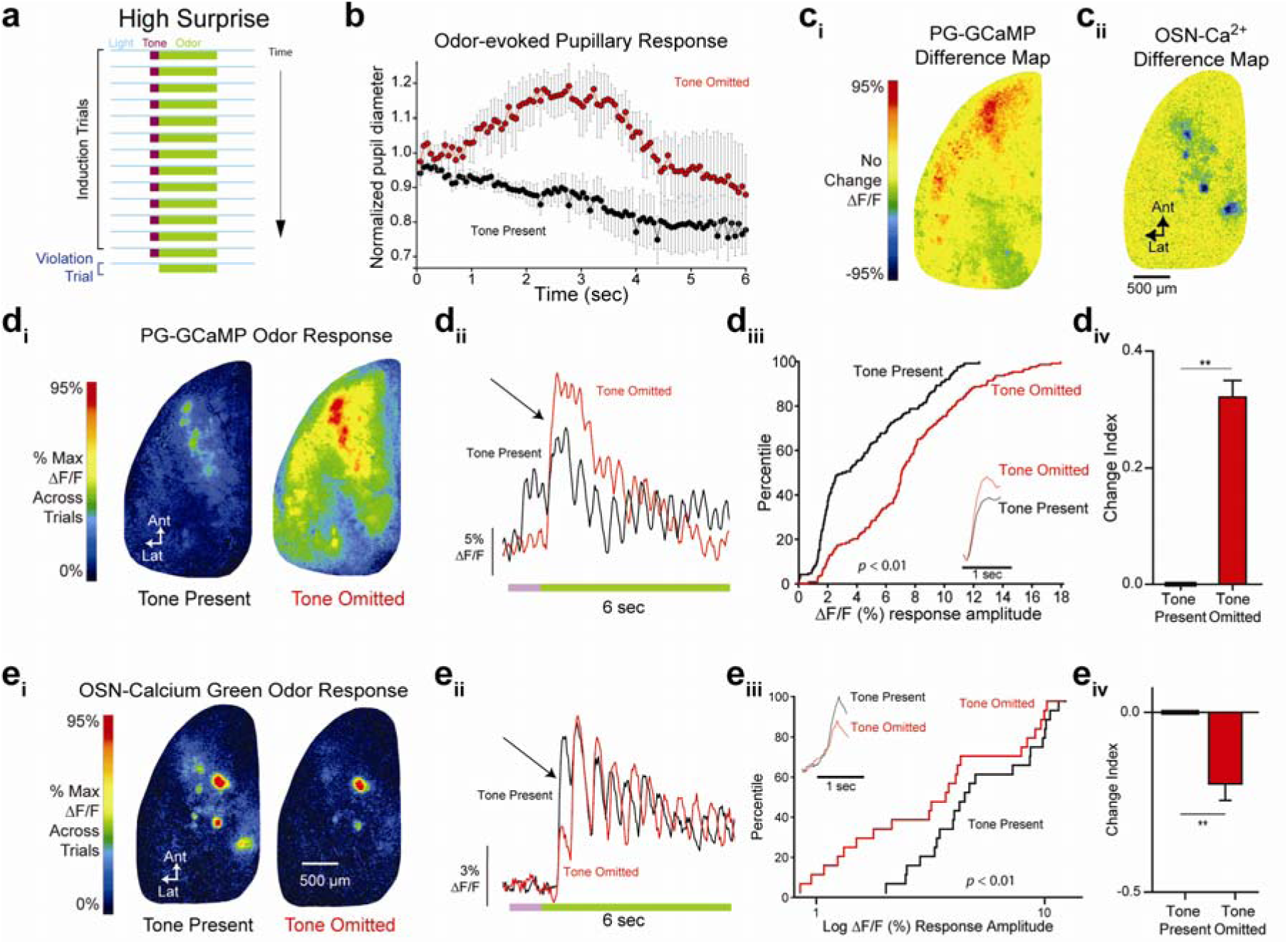
Expectation violation facilitates GABAergic signaling and suppresses presynaptic calcium in OSNs in the olfactory bulb. *A)* Paradigm schematic. *B*) Odor-evoked pupillary response shows pupil dilation on the tone omitted trial compared to the average of the previous three tone-present trials. *Cii & Ciii)* Representative pixel-by-pixel difference maps of odorant-evoked GCaMP3 (left) and Calcium Green (right) signals on the first peak of the last tone-present trial and first tone-omitted trial. Red indicates increase on the tone-omitted trial while blue indicates decrease. *Di*&*Dii*) Pseudocolor maps and fluorescence traces of odorant-evoked calcium signaling in GAD65-PG cell populations during tone-present (left/black) and tone-omitted (right/red) odorant presentations (green bar). *Diii&Div*) Odorant-evoked intraglomerular GCaMP3 signals were significantly larger on first inhalation of tone omitted (red) trials than on preceding tone-present trials (black), as shown in cumulative frequency distributions (*Diii*; Wilcoxon Signed Ranks Test *Z* = −8.56, average first sniff odorant-evoked fluorescence in inlay) and Change Index (CI) (*Div*; *t*(136) = −11.33, mean ± SE, paired t-test) across 137 glomeruli from 5 mice ** p < 0.01. *Ei&Eii*) Analogous maps and traces to *Di* & *Dii* but of presynaptic calcium signaling in OSN terminals. *Eiii&Eiv*) Odorant-evoked intraglomerular Calcium Green signals were significantly smaller on first inhalation of tone omitted (red) trials than on preceding tone-present trials (black), as shown in cumulative frequency distribution (*Eiii*; Wilcoxon Signed Ranks Test *Z* = - 3.86, average first sniff odorant-evoked fluorescence in inlay) and CI (*Eiv*; *t*(21) = 4.29, mean ± SE, paired t-test) across 22 glomeruli from 4 mice.

### Olfactory bulb interneurons respond to the omission of a learned odor

One possible function of cross-modal signaling to the olfactory bulb is to cue expectations about impending odors. To test this hypothesis, naïve Gad2-GCaMP3 mice were headfixed and presented with 13 consecutive light-tone-odor trials (Fig. 3A) at approximately one minute intervals. The light persisted for the duration of the trial with a one second tone being presented after a 3 second delay, with the odor delivered immediately upon tone termination. On the 14^th^ trial (the test trial) the light and tone were presented with same timing as on the preceding trials but the *odor was omitted* (Fig. 3A). This allowed us to observe the response evoked by cued expectation of an odor without presenting the odor itself. Remarkably, diffuse but strong *Gad2*-PG activity was observed during the period the odor was expected, with a peak response amplitude not significantly different from the preceding cued odor trial, even though no olfactory stimulus was actually presented (Fig. 3B & E). In a control group, the same paradigm was used, but odors were never presented (Fig. 3C). In these mice, endogenous *Gad2*-PG cell responses to the light and tone combination were observed (as in Fig. 1C), but they were significantly smaller than the responses to the same light and tone stimuli when they predicted an odor (Fig. 3D & E). The response magnitude difference during odor absence in these two different trial contexts suggests that cross-modally derived olfactory expectations more strongly evoke responses than repeated non-olfactory stimuli containing no information regarding future olfactory consequences. Note that in this and all subsequent experiments, the mice were not engaged in any behavioral task or receiving any reinforcement.

### Odor representations are altered in response to an omitted “warning” tone

A very pure test of expectation in sensory systems is to assess whether the *omission* of an expected cue alters the response to the stimulus the cue predicted^7^. By utilizing a sequence of predictive cues like the light-tone-odor pattern presented above, we tested the effects of omission of the odor-predictive auditory tone cue. This allowed us to observe the neural response to the same odor, presented at the same time relative to the light, with or without its directly preceding auditory cue. Naïve mice were headfixed and presented with 13 consecutive light-tone cued odor trials (as in Fig. 4A), however this expectation induction phase was followed by a 14^th^ trial in which the light and odor were presented with the same sequence and timing but the *tone was omitted* (Fig. 4A). To confirm that mice detect the change in stimulus contingency from the omission of the “warning tone” cue, we tested this paradigm in wild-type control animals while observing the mouse’s pupil. When the odor was presented without the expected tone cue, we observed a distinct dilation of the mouse’s pupil that began shortly after odor onset and was absent on the preceding three tone-present trials (Fig. 4B). When the experiment was repeated in *Gad2*-GCaMP3 mice, on the tone-omitted trial the presentation of the odor evoked a significantly larger response (Fig. 4D_i-iv_) during the first odor inhalation than on the preceding tone-present trial.

Since GABA released from *Gad2*-PG cells binds to GABA_B_ receptors on the presynaptic terminals of OSNs, suppresses N-type calcium conductance and thus neurotransmitter release^25^, we hypothesized that the increased response of *Gad2*-PG cells on the first inhalation of an odor presented without the expected tone cue would then suppress calcium influx in the OSN presynaptic terminals. To test this hypothesis, we repeated the experiment in wild-type mice whose OSN presynaptic terminals were selectively labeled with the calcium-sensitive dye Calcium Green dextran via intranasal instillation and anterograde transport^26^. These data indeed revealed a significant decrease in odor-evoked calcium flux in OSN terminals on the surprising tone-omitted trial (Fig. 4E_i-iv_). Importantly, this suppression was observed during the very first inhalation of odorant (Fig. 4E_ii_), reflecting a very rapid modulation of olfactory input. The frequency of sniff-evoked calcium transients in OSN terminals was unchanged between the tone-present and tone-omitted trials (Fig. S1). To compare the spatial element of this expectation-based modulation across the dorsal olfactory bulb, we computed difference maps between the tone-cued and tone-omitted trials for both the *Gad2*-GCaMP activity (Fig. 4C_i_), and the OSN-Calcium Green signals (Fig. 4C_ii_). Representative maps show that the *Gad*2-PG cell activity was elevated diffusely across the dorsal bulb (Fig. 4C_i_), and OSN presynaptic calcium signals were reduced in all of the glomerular foci activated by the odor (Fig. 4C_ii_). This suggests that surprising odor presentation evokes a global modulation of olfactory input across glomeruli, not only the glomeruli responsive to the cued odorant.

As a control, we repeated the tone-omission experiment with an altered paradigm in which the tone had a 50% chance of omission during baseline trials and was omitted on the 14^th^ trial as above. In wild-type mice with Calcium Green-loaded OSNs, the odor-evoked response on the 14^th^ trial (where the omission of the tone was not very surprising) was not significantly different than on the preceding tone-present trial (Fig. 5A-C). Notably, the odor responses were not different between tone-present and tone-omitted trials during the baseline trials, demonstrating that the mere presence of a tone that does not strongly correlate with odor presentation had no effect on OSN output (Fig. 5C). Finally, to confirm the importance of cross-modal input and rule out peripheral mechanisms, the tone-omission experiments were repeated in anesthetized OMP-spH mice. No effect of tone omission was observed (Fig. 5D-G), consistent with the finding that cross-modal information does not reach the PG-OSN circuit in anesthetized mice.

**Fig. 5.**
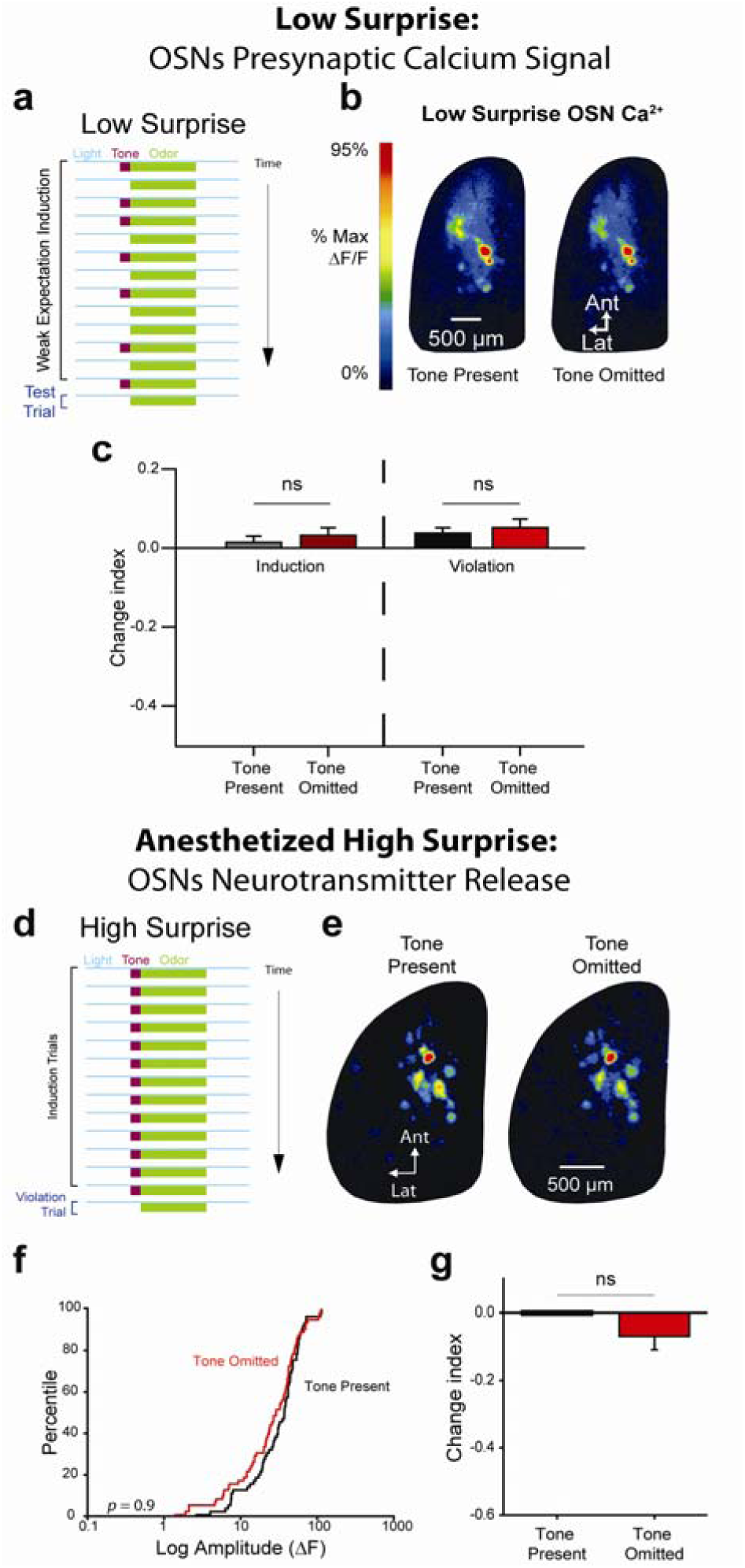
No effect of expectation when tone omission is not surprising, or in anesthetized animals. *A*) Low surprise paradigm schematic. *B*) Pseudocolor maps of odorant-evoked fluorescence on the first sniff of the last tone present trial (left) and first tone omitted trial (right). *C*) No difference (mean ± SE, paired samples t-test, *t*(19) = −0.9, *p* = 0.38); scaled JZS Bayes Factor = 3.0 in response to odorant on tone-present and tone-omitted trials (Violation). During induction phase, average of last 3 tone-present (grey) and last 3 tone-omitted (dark red) trials was not different (paired samples t-test *t*(20) = −1.01, *p* = 0.33; scaled JZS Bayes Factor = 2.8). *D*) High surprise paradigm. *E&G)* Analogous pseudocolor maps and CI measurements to *B* and *C* but in anesthetized OMP-spH mice. No difference in response amplitudes between the tone-omitted trial and preceding tone-present trial (*F*, Wilcoxon *Z* = −0.13; *G*, mean ± SE, paired samples t-test *t*(59) = 1.72, *p* = 0.09, scaled JZS Bayes Factor = 1.77, 60 glomeruli from 4 mice).

**Fig. 5.**
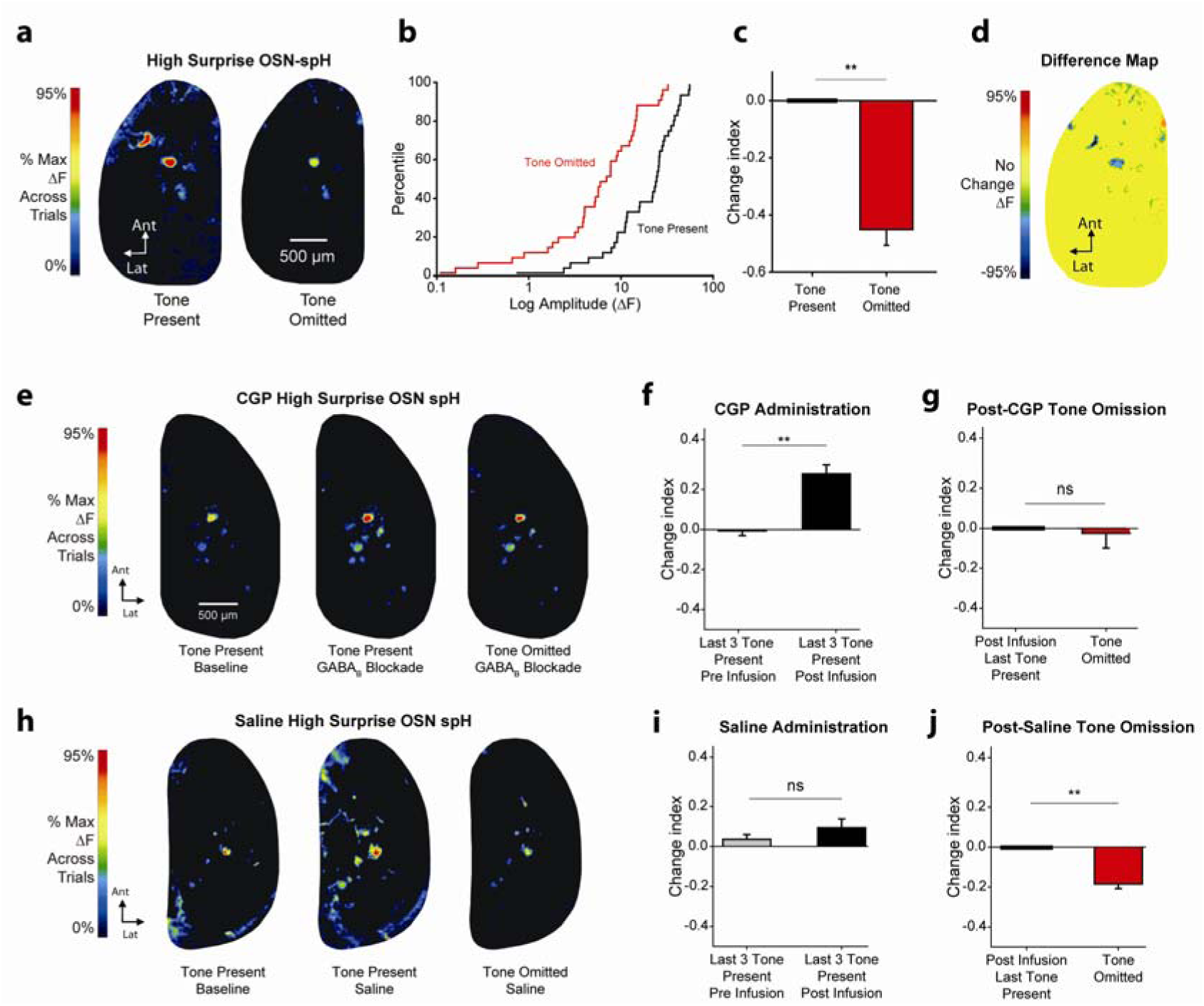
Expectation violation suppresses neurotransmitter release from OSNs via GABA_B_ receptor-mediated presynaptic inhibition. *A*) Pseudocolor maps of odorant-evoked fluorescence on last tone-present trial (left) and tone omitted trial (right) in OBs of an awake OMP-spH mouse. *B&C)* Odorant-evoked spH signals on tone-omitted (red) trials were significantly smaller than on preceding tone-present trials (black), as shown in cumulative frequency distributions (*B*; Wilcoxon Signed Ranks Test *Z* = −4.78) and Change Index (*C*; paired *t*(37) = 8.17) across 38 glomeruli from 5 mice. ** *p* < 0.01 *D*) Representative pixel-by-pixel difference map of odorant-evoked fluorescence on last tone-present trial and following tone-omitted trial. Blue indicates suppression on tone-omitted trial. *E&H*) Pseudocolor maps of odorant-evoked fluorescence on baseline tone-present trial, post-infusion tone-present trial, and post-infusion tone-omitted trial after GABA_B_ antagonist or vehicle infusion. *F&G*) GABA_B_ receptor blockade significantly enlarged odorant-evoked spH signals on last three tone present pre-infusion trials compared to last three tone present post-infusion trials (*F*; mean ± SE, paired *t*(70) = −7.61; 71 glomeruli from 4 mice). Vehicle infusion did not (*I; t*(53) = −1.84, 54 glomeruli from 5 mice; scaled JZS Bayes Factor = 1.4). *G*) After GABA_B_ receptor blockade, odorant-evoked spH signals on tone-omitted trials (red) were no different than preceding tone-present trials (black) (paired *t*(67) = −1.33; scaled JZS Bayes Factor = 3.24). *J*) After vehicle infusion, odorant-evoked spH signals on tone-omitted trial (red) were significantly smaller than on tone-present trials (black) (paired *t*(53) = 7.67).

### GABA_B_ receptor blockade blocks effect of surprise on OSNs

Presynaptic calcium flux exhibits a non-linear and temporally complex relationship to neurotransmitter release^25,27^, so we repeated the tone-omission experiment a fourth time in mice where exocytosis from OSN axon terminal populations was visualized with the fluorescent exocytosis indicator synaptopHluorin (OSN-spH mice)^28^. Consistent with the presynaptic calcium data, repeating the tone-omission experiment in OSN-spH mice revealed a significant decrease in odor-evoked neurotransmitter release from OSN terminals on the surprising, tone-omitted trial (Fig. 6A-D) compared to the preceding tone-present trial. To confirm that the information about tone-omission was being conveyed to OSNs via GABA release from OSNs, we then performed an extended version of the tone-omission experiment in a group of OMP-spH mice in which the GABA_B_ receptor antagonist CGP35348 (or vehicle control) was administered systemically midway through the baseline period. GABA_B_ receptor blockade relieved the presynaptic inhibition of OSN terminals as expected^29^, significantly enlarging the responses to the tone-cued odor compared to the pre-drug baseline (Fig. 6E & F). As hypothesized, odor presentation on a tone-omitted test trial did not evoke less neurotransmitter release from OSNs than odor presentation on the preceding tone-present trial in mice that received CGP35348 (Fig. 6E & G, right), consistent with the hypothesis that the reduced OSN synaptic output on tone-omitted trials is caused by GABA_B_ receptor activation. Control mice receiving CGP35348 infusion did exhibit expected differences in pupillary diameter on tone-omitted trials, confirming that they detected the omission of the tone (Fig. S2). OMP-spH mice that received vehicle infusion showed no significant effect of infusion and did show the expected reduction in OSN output on the tone-omitted test trial (Fig. 6H-J). While it may seem counterintuitive for the olfactory system to suppress OSN neurotransmitter release when an odor is presented unexpectedly, in fact the suppression of glutamate release via presynaptic GABA_B_ receptors can potentially help prevent neurotransmitter depletion and AMPA receptor desensitization, thus increasing the synapses’ ability to sustain activity during periods of strong stimulation^12,27,30,31^.

## Discussion

Most models of sensory processing presume that brain regions tasked with early sensory processing are limited to extracting the sensory inputs to their single sensory modality and presenting that “bottom-up” information to higher brain regions for interpretation and contextualization. However, the present data prove that in awake animals, sensory processing as early as the olfactory nerve can be influenced by odor-predictive cues even if they originate in non-olfactory modalities. Even some of the first neurons to receive afferent olfactory signals from the periphery are also responsive to infrequent visual, auditory, and somatosensory stimuli. Cross-modal sensory cueing may thus play a more fundamental role than previously imagined.^32-34^ Surprisingly, these modulations occurred even though the animal was never engaged in an explicit behavioral task. It remains to be determined what underlying anatomical projections convey this information, what level of detail is communicated, and whether attentional mechanisms^35,36^ play a role in this integration as well. It is conceivable that the nature of an expectation (eg passive or task-specific) engages different cortical and neuromodulatory areas that differentially engage olfactory bulb circuits. None-the-less, the modulation of OSN activity by cognitive factors like expectations and cross-modal stimulus contingencies suggests that there is actually no such thing as a purely bottom-up sensory representation in the olfactory system and that primary sensory areas may not be as dedicated as previously believed.

## Acknowledgements

This work was supported by NIH grants DC009442, MH101293, and DC013090 to JPM. LAC and JPM conceived and designed the experiments. LAC, AHM, CDF, DJT and JPM collected and analyzed the data. JPM and LAC prepared the manuscript. All data reported here are available upon request. We thank Randy Gallistel, Kasia Bieszczad, Mimi Phan, and David Vicario for helpful comments.

## Materials and Methods

### Subjects

A total of 73 mice were used in these experiments of which 36 were male and 37 were female. All mice were between 2 and 10 months of age the time of experimentation. The fluorescent calcium indicator GCaMP3^40^ of GCaMP6f was expressed in cells expressing the GAD65-encoding gene *gad2* via cre recombinase-mediated excision of a floxed STOP codon^24,41^. PG-GCaMP mice were F1 crosses of the Gad2^tm2(cre)Zjh^/6 (Jackson Labs stock #010802)^42^ with either the B6;129S-Gt(ROSA)26Sor^tm38(CAG-GCaMP3)Hze^/6 line (Jackson Labs stock #014538)^43^ or B6;129S-Gt(ROSA)26Sor^tm95.1(CAG-GCaMP6 f)Hze^/J line (Jackson Labs stock #024105), except for comparison mice (Fig. 2D) where GCaMP6f was introduced via viral vector (see below). OMP-spH mice were heterozygous for olfactory marker protein and spH on a mixed albino C57BL/6 and 129 ^44 44 44^ background as previously reported^44,45^. Wild-type mice for calcium imaging experiments were 129 strain mice to facilitate comparison to the OMP-spH mice. All mice were maintained on a 12:12 h light:dark cycle and given *ad libitum* food and water. All experiments were conducted in accordance with protocols approved by the Rutgers University Institutional Animal Care and Use Committee.

### Restraint training

All mice underwent 2 to 4 daily 60-minute restraint training sessions in custom ventilated restraint tubes ∼ 2.8cm in diameter prior to surgery. Following headcap implantation, some mice received up to two additional 90 minute restraint sessions in which they were secured to a custom headholder via the implanted headcap in a mockup of the imaging apparatus to habituate them to restraint.

### Headcap and cranial window implantation

Headcaps and cranial windows were implanted as previously reported^45,46^. Briefly, mice were anesthetized with 1000 µg/kg dexmedetomadine (Dexdomitor, Orion Corporation) and 70 mg/kg ketamine administered i.p. with bupivicaine (∼ 0.25 mL at 0.25%, s.c.) as a local anesthetic at the incision site. The scalp was shaved, washed with three cycles of Betadine and 70% ethanol and then surgically opened with a midline incision. The periosteal membrane was removed and the skull dried with a 70% ethanol solution. A custom acrylic headcap was secured using cyanoacrylate and dental acrylic and the skull overlying both olfactory bulbs was thinned to transparency and coated with a thin layer of cyanoacrylate. The wound margins were secured with tissue adhesive (Vetbond, 3M) and the window covered by a protective metal shield secured to the headcap. For calcium imaging OSNs were loaded with 4 µL 4% Calcium Green dextran potassium salt, a dextran-conjugated, fluorescent calcium-sensitive dye (Life Technologies, Grand Island, NY) by intranasal instillation during headcap implantation after the manner of Wachowiak and Cohen^26^. Briefly, a microloader attached to a Hamilton syringe was inserted 7mm into each naris. The animal was rotated onto its back and 2 µL 2% Triton was instilled initially followed by 2 µL 4% Calcium Green after 1 min. The remaining 2µL was instilled 5 min later. After an additional 5 min the animal was rotated onto its side to allow the dye to reach all areas of the epithelium. The same procedure was performed on the contralateral side after an additional 15 min. Anesthesia was reversed with 1 mg/kg s.c. atipamezole (Antisedan, Orion Corporation). Mice were given 24-48 hours recovery time prior to further experimentation. All animals were singly housed following headcap implantation.

### Viral vector instillation

Mice were anesthetized with 1000 µg/kg dexmedetomadine (Dexdomitor, Orion Corporation) and 70 mg/kg ketamine administered i.p. with bupivicaine (∼ 0.25 mL at 0.25%, s.c.) as a local anesthetic at the incision site. The scalp was shaved, washed with three cycles of Betadine and 70% ethanol and then surgically opened with a midline incision. The periosteal membrane was removed and the skull dried with a 70% ethanol solution. An FG 1/2 burr was used to make a hole in the skull over each olfactory bulb. A gastight 1.0µL Hamilton syringe (1700 series; Hamilton Company) was used to infuse (Quintessential Stereotaxic Injector, Stolting) 0.5µL of AAV1.Syn.Flex.GCaMP6f.WPRE.SV40 (University of Pennyslvania Vector Core) into the glomerular layer at a rate of 0.1µL/minute. After infusion completion, the syringe remained in the tissue for five additional minutes to allow for diffusion. The process was then repeated for the contralateral olfactory bulb. The midline incision was closed and secured with either tissue adhesive (Vetbond, 3M) or interrupted 6.0 silicone coated braided sutures (Sofsilk, Covidien).

### Optical imaging of olfactory bulb function

For optical imaging, the protective shield overlying the cranial window was removed and the mouse was secured to the headholder. A drop of Ringer’s solution (140 mM NaCl, 5 mM KCl, 1mM CaCl_2_, 1mM MgCl_2_, 10mM HEPES and 10mM dextrose) and a glass coverslip was positioned over the cranial window to ensure a flat optical surface. Optical imaging was performed as previously described^46-48^. Briefly, the dorsal surface of both olfactory bulbs was visualized using a custom imaging apparatus including a 4x, 0.28 NA Olympus macro objective lens and images were acquired at 7 Hz (spH) or 25 Hz (Calcium Green or GCaMP) using a low-light, back-illuminated CCD camera with a resolution of 256×256 pixels (RedShirtImaging, Decatur, GA). Epiillumination was provided by a 488-nm wavelength LED from ThorLabs (Newton, NJ). The odorant methyl valerate was presented in 6-sec trials at a concentration of ∼ 2% saturated vapor via vapor-dilution olfactometer using nitrogen as the carrier.

### Pupillometry

Mice for pupillometry experiments were restraint trained and underwent cranial window implantation as above. Mice were head-fixed and positioned under a resonance-scanned (Neurolabware, Los Angeles, CA) titanium:sapphire ultrafast pulsed laser (Coherent Chameleon Ultra II) tuned to 920 nm (infrared). Images of the pupil were captured at 15.5 frames per second using a GigE camera sensitive to infrared light. IR light passed through the brain, transmitted through the orbit, and was emitted from the pupil. Pupil area was measured in each frame using custom software in Matlab based on the size of the circle that best fit the infrared light escaping the pupil. To mimic the illumination light during wide-field imaging, each pupillometry trial included illuminating the mouse with fiber optic-coupled blue light from a 470 nm LED identical to that used in the wide-field imaging apparatus (Thor Labs, Newton, NJ). Auditory stimuli were matched for position and sound pressure level. Two mice were excluded from analysis because the pupil was frequently obscured by partial closure of the lower eyelid, but exhibited qualitatively similar behavior to those reported here. An additional mouse was excluded because its pupil was completely dilated and exhibited no pupillary responses to any stimulus, suggesting a ceiling effect. To improve signal-to-noise ratio, the average of the final three tone-present trials was calculated as a baseline for each mouse. Data were normalized for averaging across mice.

### Tracheotomy

Prior to surgery, a borosilicate glass capillary tube (1B120F-4; World Precision) was heated, bent into an S-shape, and pulled and PE-50 tubing (Scientific Commodities, Inc) was pulled and secured onto the glass capillary tube in order to extend the opening of the capillary tube away from the thoracic cavity. Following cranial window implantation, mice were placed on their back and given bupivicane along the median cervical skin. This area was shaved and cleaned with alternating washes of 70% ethanol solution and Betadine surgical scrub. A single incision was made along this axis. The muscle was retracted to expose the trachea. An incision was made in the sublaryngeal region and the glass tube was inserted into the trachea with the open end facing the lungs and the other end outside the body. The tube was secured with suture that held the trachea tightly around the tube. The skin around the tracheal tube was then closed and secured with tissue adhesive (Vetbond, 3M). Animals were then secured to the head-holder and allowed to wake up in the widefield imaging apparatus. Mice breathed freely through the tracheal tube, with no airflow possible through the nasal passages. The tracheal tube was monitored for condensation and cleared when necessary.

### Drug administration via intraperitoneal cannula

Mice were randomly assigned to receive either saline or CGP35348 infusions. Animals were lightly anesthetized with isoflurane (Aerrane, Baxter Healthcare, Deerfield, IL) and implanted with a pre-loaded length of polyethylene tubing (Intramedic PE 10, Becton Dickinson) via an 18G guide needle, attached to a 1.0 mL syringe. CGP35348 or saline was administered via intraperitoneal cannula after 9 to 11 tone-present Expectation Induction trials were presented to collect a stable baseline. After drug infusion, Expectation Induction trials continued until OSN response amplitudes plateaued (range: 11 to 17 trials, with number of trials for control animals yoked to CGP-infused animals), at which point the tone-omitted Expectation Violation trial was administered.

### Optical neurophysiology data analysis

Putative glomeruli were hand-selected based on their odorant-evoked change in fluorescence and their responses. For spH data, which produces an integrative signal, the response of each glomerulus was measured as the peak fluorescence within 4 sec of the odorant offset minus the pre-odor baseline fluorescence on each trial. For calcium data, inhalation-induced peaks in the fluorescence signal were measured relative to the pre-peak trough and normalized to the baseline fluorescence, producing a ΔF/F measure for each inhalation. Trials with movement artifacts were discarded. Glomerular responses were pooled across mice where indicated. For multimodal and odor omission experiments, whole bulb ROIs were selected to capture diffuse non-glomerular signals. The experimenter analyzing data was blind to trial type until response amplitude quantification was complete.

To facilitate comparisons across experiments, we expressed population averages in terms of a change index, defined as (test trial - reference trial)/(test trial + reference trial), after the manner of Kato et al.^49^ This index equals −1 if the response on the test trial is completely abolished, equals 0 if the response on the test and reference trials are the same size, and asymptotically approaches 1 as the response on the test trials exceeds the reference trial. This approach is superior to traditional “percent change” approaches because it prevents unidirectional bounding, i.e. that responses can be more than 100% increased but cannot be more than 100% decreased.

Pilot experiments were used to assess approximate power utilizing these within-subjects designs. We have generally presented both normalized group data (with appropriate parametric statistics) as well as distributions (and appropriate non-parametric statistics of raw values) of all data points that went into the parametric group averages. Paired samples t-tests were used for planned comparisons within-subjects between two trial types. Wilcoxon Signed Rank tests were used for analyses of related distributions. Mixed model ANOVAs were used for designs with within and between subjects manipulations except for modality comparisons in figure 1. In this case, all subjects did not receive all stimuli, thus not allowing for complete matched cases. Instead, a Mixed Linear Model was used. All tests were two-tailed. One glomerulus (out of 21) was included in analysis in figure 6c (left) that was not included in figure 6c (right). The left portion of the graph are 3 trial averages, where as the right portion of the graph is single trial comparisons and was missing from one of the single trials. Three glomeruli (out of 71) were included in figure 5f, but not 5g for the same reason.

### Comparison of GCaMP3 expression to OSN afferents (Fig. 1B)

OSNs were labeled with AlexaFluor 568-conjugated dextran (Life Technologies) via intranasal infusion as described above. GCaMP3 was expressed in via cre recombinase-mediated excision of a floxed STOP codon in cells expressing cre recombinase from the *gad2* locus as described in the main text. Structural images were captured through a cranial window in anesthetized mice using a 2-photon microscope (Neurolabware) with a 25X objective with a 1.05 numerical aperture (Nikon).

## References

1. Zelano, C., Mohanty, A. & Gottfried, J. A. Olfactory predictive codes and stimulus templates in piriform cortex. Neuron 72, 178–187, doi:10.1016/j.neuron.2011.08.010 (2011).

2. Gottfried, J. A. & Dolan, R. J. The nose smells what the eye sees: crossmodal visual facilitation of human olfactory perception. Neuron 39, 375–386 (2003).

3. Naatanen, R., Kujala, T. & Winkler, I. Auditory processing that leads to conscious perception: a unique window to central auditory processing opened by the mismatch negativity and related responses. Psychophysiology 48, 4–22, doi:10.1111/j.1469-8986.2010.01114.x (2011).

4. den Ouden, H. E., Friston, K. J., Daw, N. D., McIntosh, A. R. & Stephan, K. E. A dual role for prediction error in associative learning. Cerebral cortex 19, 1175–1185, doi:10.1093/cercor/bhn161 (2009).

5. Egner, T., Monti, J. M. & Summerfield, C. Expectation and surprise determine neural population responses in the ventral visual stream. The Journal of neuroscience: the official journal of the Society for Neuroscience 30, 16601–16608, doi:10.1523/JNEUROSCI.2770-10.2010 (2010).

6. Langner, R. et al. Modality-specific perceptual expectations selectively modulate baseline activity in auditory, somatosensory, and visual cortices. Cerebral cortex 21, 2850–2862, doi:10.1093/cercor/bhr083 (2011).

7. Sanmiguel, I., Saupe, K. & Schroger, E. I know what is missing here: electrophysiological prediction error signals elicited by omissions of predicted “what” but not “when”. Frontiers in human neuroscience 7, 407, doi:10.3389/fnhum.2013.00407 (2013).

8. Gill, P., Woolley, S. M., Fremouw, T. & Theunissen, F. E. What’s that sound? Auditory area CLM encodes stimulus surprise, not intensity or intensity changes. Journal of neurophysiology 99, 2809–2820, doi:10.1152/jn.01270.2007 (2008).

9. Allen, W. E. et al. Global Representations of Goal-Directed Behavior in Distinct Cell Types of Mouse Neocortex. Neuron 94, 891–907 e896, doi:10.1016/j.neuron.2017.04.017 (2017).

10. Shao, Z., Puche, A. C., Kiyokage, E., Szabo, G. & Shipley, M. T. Two GABAergic intraglomerular circuits differentially regulate tonic and phasic presynaptic inhibition of olfactory nerve terminals. Journal of neurophysiology 101, 1988–2001, doi:10.1152/jn.91116.2008 (2009).

11. Aroniadou-Anderjaska, V., Zhou, F. M., Priest, C. A., Ennis, M. & Shipley, M. T. Tonic and synaptically evoked presynaptic inhibition of sensory input to the rat olfactory bulb via GABA(B) heteroreceptors. Journal of neurophysiology 84, 1194–1203 (2000).

12. McGann, J. P. Presynaptic inhibition of olfactory sensory neurons: new mechanisms and potential functions. Chemical senses 38, 459–474, doi:10.1093/chemse/bjt018 (2013).

13. Murphy, G. J., Darcy, D. P. & Isaacson, J. S. Intraglomerular inhibition: signaling mechanisms of an olfactory microcircuit. Nature neuroscience 8, 354–364, doi:10.1038/nn1403 (2005).

14. Markopoulos, F., Rokni, D., Gire, D. H. & Murthy, V. N. Functional properties of cortical feedback projections to the olfactory bulb. Neuron 76, 1175–1188, doi:10.1016/j.neuron.2012.10.028 (2012).

15. Petzold, G. C., Hagiwara, A. & Murthy, V. N. Serotonergic modulation of odor input to the mammalian olfactory bulb. Nature neuroscience 12, 784–791, doi:10.1038/nn.2335 (2009).

16. D’Souza, R. D. & Vijayaraghavan, S. Nicotinic receptor-mediated filtering of mitral cell responses to olfactory nerve inputs involves the alpha3beta4 subtype. The Journal of neuroscience: the official journal of the Society for Neuroscience 32, 3261–3266, doi:10.1523/JNEUROSCI.5024-11.2012 (2012).

17. Liu, S. et al. Muscarinic receptors modulate dendrodendritic inhibitory synapses to sculpt glomerular output. The Journal of neuroscience: the official journal of the Society for Neuroscience 35, 5680–5692, doi:10.1523/JNEUROSCI.4953-14.2015 (2015).

18. Rothermel, M., Carey, R. M., Puche, A., Shipley, M. T. & Wachowiak, M. Cholinergic inputs from Basal forebrain add an excitatory bias to odor coding in the olfactory bulb. The Journal of neuroscience: the official journal of the Society for Neuroscience 34, 4654–4664, doi:10.1523/JNEUROSCI.5026-13.2014 (2014).

19. Steinfeld, R., Herb, J. T., Sprengel, R., Schaefer, A. T. & Fukunaga, I. Divergent innervation of the olfactory bulb by distinct raphe nuclei. The Journal of comparative neurology 523, 805–813, doi:10.1002/cne.23713 (2015).

20. Eckmeier, D. & Shea, S. D. Noradrenergic plasticity of olfactory sensory neuron inputs to the main olfactory bulb. The Journal of neuroscience: the official journal of the Society for Neuroscience 34, 15234–15243, doi:10.1523/JNEUROSCI.0551-14.2014 (2014).

21. Suzuki, Y., Kiyokage, E., Sohn, J., Hioki, H. & Toida, K. Structural basis for serotonergic regulation of neural circuits in the mouse olfactory bulb. The Journal of comparative neurology 523, 262–280, doi:10.1002/cne.23680 (2015).

22. Fast, C. D. & McGann, J. P. Amygdalar gating of early sensory processing through interactions with locus coeruleus. Journal of Neuroscience 37, 3085–3101 (2017).

23. Park, S. & McGann, J. P. Functional inactivation of the anterior olfactory nucleus enhances odor-evoked periglomerular cell activity in the mouse olfactory bulb. (In Review).

24. Madisen, L. et al. Transgenic mice for intersectional targeting of neural sensors and effectors with high specificity and performance. Neuron 85, 942–958, doi:10.1016/j.neuron.2015.02.022 (2015).

25. Wachowiak, M. et al. Inhibition of olfactory receptor neuron input to olfactory bulb glomeruli mediated by suppression of presynaptic calcium influx. Journal of neurophysiology 94, 2700–2712, doi:10.1152/jn.00286.2005 (2005).

26. Wachowiak, M. & Cohen, L. B. Representation of odorants by receptor neuron input to the mouse olfactory bulb. Neuron 32, 723–735 (2001).

27. Murphy, G. J., Glickfeld, L. L., Balsen, Z. & Isaacson, J. S. Sensory neuron signaling to the brain: properties of transmitter release from olfactory nerve terminals. The Journal of neuroscience: the official journal of the Society for Neuroscience 24, 3023–3030, doi:10.1523/JNEUROSCI.5745-03.2004 (2004).

28. Bozza, T., McGann, J. P., Mombaerts, P. & Wachowiak, M. In vivo imaging of neuronal activity by targeted expression of a genetically encoded probe in the mouse. Neuron 42, 9–21 (2004).

29. McGann, J. P. et al. Odorant representations are modulated by intra-but not interglomerular presynaptic inhibition of olfactory sensory neurons. Neuron 48, 1039–1053, doi:10.1016/j.neuron.2005.10.031 (2005).

30. Brenowitz, S., David, J. & Trussell, L. Enhancement of synaptic efficacy by presynaptic GABA(B) receptors. Neuron 20, 135–141 (1998).

31. Brenowitz, S. & Trussell, L. O. Minimizing synaptic depression by control of release probability. The Journal of neuroscience: the official journal of the Society for Neuroscience 21, 1857–1867 (2001).

32. van den Brink, R. L. et al. Subcortical, Modality-Specific Pathways Contribute to Multisensory Processing in Humans. Cerebral cortex 24, 2169–2177, doi:DOI 10.1093/cercor/bht069 (2014).

33. Wu, C., Stefanescu, R. A., Martel, D. T. & Shore, S. E. Listening to another sense: somatosensory integration in the auditory system. Cell and tissue research, doi:10.1007/s00441-014-2074-7 (2014).

34. Spence, C. Multisensory Flavor Perception. Cell 161, 24–35, doi:10.1016/j.cell.2015.03.007 (2015).

35. Li, G. & Cleland, T. A. A two-layer biophysical model of cholinergic neuromodulation in olfactory bulb. The Journal of neuroscience: the official journal of the Society for Neuroscience 33, 3037–3058, doi:10.1523/JNEUROSCI.2831-12.2013 (2013).

36. D’Souza, R. D. & Vijayaraghavan, S. Paying attention to smell: cholinergic signaling in the olfactory bulb. Frontiers in synaptic neuroscience 6, 21, doi:10.3389/fnsyn.2014.00021 (2014).

37. Verhagen, J. V., Wesson, D. W., Netoff, T. I., White, J. A. & Wachowiak, M. Sniffing controls an adaptive filter of sensory input to the olfactory bulb. Nature neuroscience 10, 631–639, doi:10.1038/nn1892 (2007).

38. Cenier, T., McGann, J. P., Tsuno, Y., Verhagen, J. V. & Wachowiak, M. Testing the sorption hypothesis in olfaction: a limited role for sniff strength in shaping primary odor representations during behavior. The Journal of neuroscience: the official journal of the Society for Neuroscience 33, 79–92, doi:10.1523/JNEUROSCI.4101-12.2013 (2013).

39. Wesson, D. W., Donahou, T. N., Johnson, M. O. & Wachowiak, M. Sniffing behavior of mice during performance in odor-guided tasks. Chemical senses 33, 581–596, doi:10.1093/chemse/bjn029 (2008).

40. Akerboom, J. et al. Optimization of a GCaMP calcium indicator for neural activity imaging. The Journal of neuroscience: the official journal of the Society for Neuroscience 32, 13819–13840, doi:10.1523/JNEUROSCI.2601-12.2012 (2012).

41. Wachowiak, M. et al. Optical dissection of odor information processing in vivo using GCaMPs expressed in specified cell types of the olfactory bulb. The Journal of neuroscience: the official journal of the Society for Neuroscience 33, 5285–5300, doi:10.1523/JNEUROSCI.4824-12.2013 (2013).

42. Taniguchi, H. et al. A resource of Cre driver lines for genetic targeting of GABAergic neurons in cerebral cortex. Neuron 71, 995–1013, doi:10.1016/j.neuron.2011.07.026 (2011).

43. Zariwala, H. A. et al. A Cre-dependent GCaMP3 reporter mouse for neuronal imaging in vivo. The Journal of neuroscience: the official journal of the Society for Neuroscience 32, 3131–3141, doi:10.1523/JNEUROSCI.4469-11.2012 (2012).

44. Czarnecki, L. A. et al. In vivo visualization of olfactory pathophysiology induced by intranasal cadmium instillation in mice. Neurotoxicology 32, 441–449, doi:10.1016/j.neuro.2011.03.007 (2011).

45. Czarnecki, L. A. et al. Functional rehabilitation of cadmium-induced neurotoxicity despite persistent peripheral pathophysiology in the olfactory system. Toxicological sciences: an official journal of the Society of Toxicology 126, 534–544, doi:10.1093/toxsci/kfs030 (2012).

46. Kass, M. D., Pottackal, J., Turkel, D. J. & McGann, J. P. Changes in the neural representation of odorants after olfactory deprivation in the adult mouse olfactory bulb. Chemical senses 38, 77–89, doi:10.1093/chemse/bjs081 (2013).

47. Kass, M. D., Moberly, A. H., Rosenthal, M. C., Guang, S. A. & McGann, J. P. Odor-specific, olfactory marker protein-mediated sparsening of primary olfactory input to the brain after odor exposure. The Journal of neuroscience: the official journal of the Society for Neuroscience 33, 6594–6602, doi:10.1523/JNEUROSCI.1442-12.2013 (2013).

48. Kass, M. D., Rosenthal, M. C., Pottackal, J. & McGann, J. P. Fear learning enhances neural responses to threat-predictive sensory stimuli. Science 342, 1389–1392, doi:10.1126/science.1244916 (2013).

49. Kato, H. K., Chu, M. W., Isaacson, J. S. & Komiyama, T. Dynamic sensory representations in the olfactory bulb: modulation by wakefulness and experience. Neuron 76, 962–975, doi:10.1016/j.neuron.2012.09.037 (2012).

