## Supplementary Materials for "Multisensory expectations shape olfactory input to the brain"

**Supplementary Analyses**

*No Contribution of Odor Sampling Was Observed*

The present results cannot be directly attributed to changes in odor sampling behavior^37^. In mice (and unlike our work in rats^38^), high-frequency investigatory sniffing is observed in some odor-guided behavioral tasks but not others and is more strongly predicted by reward availability than by odor presentation^39^. Analysis of sniff-locked OSN calcium transients revealed no difference in sniffing frequency during the odor presentation on tone-omitted and tone-present trials in the High Surprise paradigm (Supplementary Fig. 1) despite the clear difference in OSN synaptic output during that time (Fig. 4A-D). Similarly, no differences in sniffing frequency were observed between tone-present and tone-absent trials on the Low Surprise paradigm (Fig. 5A-C). Moreover, the abolition of the effects of expectation violation on OSNs by GABA_B_ receptor blockade (Fig. 4E-G) is more consistent with a local circuit effect than a behavioral change.

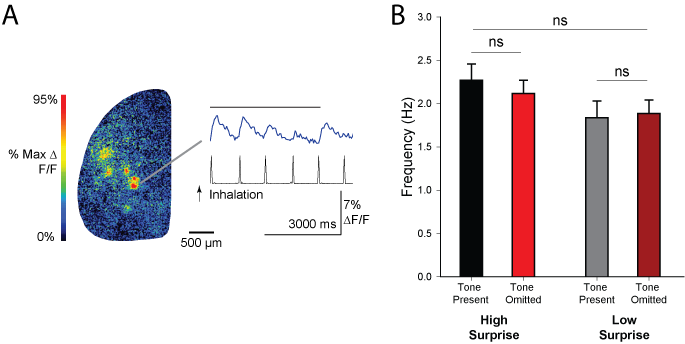

**Supplementary** **Figure 1. Expectation alters OSN response patterns without a change in odor sampling.** *A*) Pseudocolor map and example fluorescence (blue, top) and respiration (black, lower) traces from an awake mouse during an odor presentation (black bar) showing that Calcium Green signal peaks phase-lock to inhalations. *B*) Summary data (mean ± SE) showing inhalation frequency (inferred from OSN-Calcium Green signal peaks) during the high and low surprise experiments. Mixed-model ANOVA revealed no change in sampling frequency between last tone-present trial and first tone-omitted trial (*F*(1, 40) = 0.41, *p* = 0.52) in either paradigm, nor between experimental paradigms (*F*(1, 40) = 2.08, *p* = 0.16) and no significant interaction (*F*(1, 40) = 1.54, *p* = 0.22).

*Expectation Violations Were Detected Despite CGP34348 Administration*

In Fig. 4, the High Surprise Paradigm was lengthened to permit drug administration during the expectation induction phase, and the expectation violation trial was presented only after the administration of GABA_B_ receptor antagonist CGP35348 or vehicle control. To confirm that the mice were still able to detect the violation after CGP35348 administration, we repeated the CGP-administration experiment in 5 wild-type mice while monitoring pupil diameter as in Fig. 3B). As in mice undergoing the standard High Surprise paradigm, there was indeed a significantly larger pupil dilation on the tone-omitted trial than on the preceding tone-present trials (Supplementary Fig. 2).

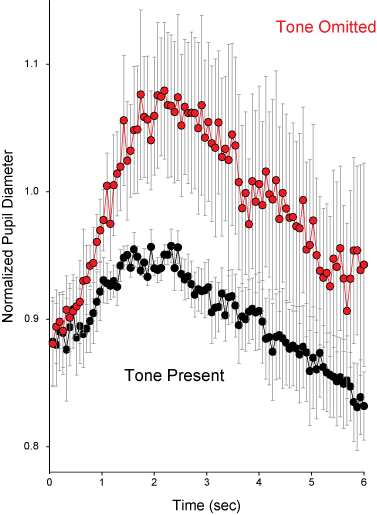

**Supplementary Figure 2. Expectation violation induces pupil dilation under GABA_B_ blockade.** Administration of GABA_B_ antagonist CGP35348 did not prevent the expression of pupil dilation after odor presentation without the expected warning tone (red). This dilation was larger than on preceding tone present trials (black; average of three preceding trials).
